# The contribution of melanization to *Drosophila* survival changes with *Enterococcus faecalis* V583 genomic content

**DOI:** 10.1101/329144

**Authors:** Neuza Teixeira, António Jacinto, Maria de Fátima Silva Lopes

**Author notes:** Corresponding author: IBET, Quinta do Marquês, Estação Agronómica Nacional,Apartado 12, 2781-901 Oeiras, Portugal.

## Abstract

*Enterococcus faecalis* is a human opportunist pathogen able to infect and kill *Drosophila*. Previous studies proved that *E. faecalis* carrying the Fsr quorum sensing system are extremely virulent. Fsr is the regulator of two important virulence factors, gelatinase and serine protease, which cause death of *Drosophila* adult flies by decreasing its tolerance to infection. The exact mechanism underlying the toxicity of these *E. faecalis* virulence factors is nevertheless not known, in particular the way they interfere with the host immune response. In the present study, we investigated the influence of Fsr-GelE-SprE bacterial factors on different immunity responses, namely antimicrobial peptide production, phagocytosis and melanization. Using *E. faecalis* V583 wild type and *E. faecalis* V583 Δ*fsrB*Δ*gelE*Δ*sprE* mutant we showed that both drosomycin production and phagocytosis were activated to similar levels by the two bacterial strains. However, fly pupae infected with the mutant strain showed less melanization and higher survival rates when compared to pupae infected with wild type bacteria. Using adult flies carrying the *PPO1*^*Δ*^ *PPO2*^*Δ*^ mutation, we found that absence of melanization had a different impact in survival of the flies when infected with the two *E. faecalis* strains. *PPO1*^*Δ*^, *PPO2*^*Δ*^ mutant flies were more tolerant to *E. faecalis* deprived of its major virulence factors. By showing that the presence of the *E. faecalis* proteases completely alters the impact of melanization activation on *Drosophila* tolerance, this study provides new clues on the interactions between *E. faecalis* virulence factors and the fly´s immune system. Future studies on *Drosophila* immunity should consider the pathogen genomic content.

## INTRODUCTION

In order to cause disease and death, pathogens must overcome the host´s immune defenses. Understanding how the host immune defense mechanisms react to pathogens and how pathogens inflict disease on the host can therefore provide us with clues to fight those more efficiently. Among the most challenging pathogens are the opportunistic ones, namely *Enterococcus faecalis*, which are commensal to humans but can cause disease in patients with impaired immune systems. Enterococci are natural inhabitants of the oral cavity, intestinal tract and female genital tract of both human and animals. In contrast to the beneficial role they play in intestinal homeostasis, these organisms are becoming increasingly important to human health as leading causes of nosocomial infections. They are prevalent in the nosocomial environment, causing infections of the urinary tract, bloodstream, intra-abdominal and pelvic regions, surgical sites and central nervous system (1). To do so, they rely on several mechanisms including the *fsr* operon in the case of *E. faecalis*. The *fsr* (*<u>E</u>nterococcus faecalis* <u>r</u>egulator) two component system, a homologue of the *agr* system in *Staphylococcus aureus*, is a quorum sensing-dependent regulatory system that regulates the expression of two other important virulence factors, *gelE* and *sprE*. These genes encode, respectively, gelatinase (GelE), an extracellular zinc metalloprotease, and SprE, a serine protease (2-4).

Recently, our Lab provided evidence for their role, and also for Fsr function, in *Drosophila melanogaster* mortality (5). *D. melanogaster* (fruit fly) is a powerful model organism to understand both the molecular mechanisms regulating the activation of innate immune response and to screen for bacterial effectors involved in virulence (6). The fruit fly has a multilayered immune system consisting of at least seven defensive mechanisms: regulation of the native microbiota in the gut through antimicrobial peptides (AMPs) and reactive oxygen species; the barrier epithelial response, which recognizes infections and wounds, produces local AMPs and sends signals to the rest of the body; the clotting response, which not only seals wounds and prevents bleeding, but can physically trap bacteria; the phenoloxidase response, which deposits melanin at the site of an immune reaction, releasing potentially antimicrobial reactive oxygen species; the phagocytic response, through which phagocytes can kill microbes directly by either encapsulation or phagocytosis, or indirectly by releasing systemic signals; the systemic AMP response, which involves the release of massive quantities of AMPs from the fat body (the liver analog) into the circulation (7); and the RNAi response, which is required to fight viral infections.

The expression of AMPs, regulated by the Toll and Imd pathways (8), can take a few hours to a few days to occur. In contrast, a more immediate immune response, induced within a few minutes after infection, is melanization (9). This is considered to be the earliest and most acute reaction of insects against pathogens upon injury (9) and is used to encapsulate and sequester pathogens too large to be phagocytized (10). During melanization reaction, phenols are oxidized to quinolones, which then polymerize to form melanin that is deposited around intruding microorganisms to help sequester them at the wound site. The quinolone substances and other reactive oxygen intermediates are thought to be directly toxic to microorganisms. Melanin synthesis is the final product of the proteolytic cascade leading to the cleavage of prophenoloxidase (proPO) to phenoloxidase (PO).

We have shown that the *E. faecalis* virulence factors Fsr, GelE and SprE are necessary to cause *Drosophila* mortality upon infection (5). However, it remains unclear how these factors control this process. In the present study we asked how different aspects of the immune response in *Drosophila* were affected upon infection with *E. faecalis* and how that depends on the Fsr, GelE and SprE machinery. We found that important resistance mechanisms, such as drosomycin expression and phagocytosis, were not altered in the absence of Fsr and the proteases. In contrast, the melanization response was severely affected in flies infected with wild type but not with Fsr mutant bacteria. Furthermore, we show that outcome of the impairment in the melanization reaction in infected flies depends on the genomic content of the infecting *E. faecalis* strain.

## MATERIALS AND METHODS

### Bacterial Strains

Strains used in this study are listed in Table 1. Enterococcal strains were grown in BHI (Brain Heart Infusion) medium at 37°C, and *Micrococcus luteus* strain was grown in LB medium at 37°C with agitation.

**Table 1.**
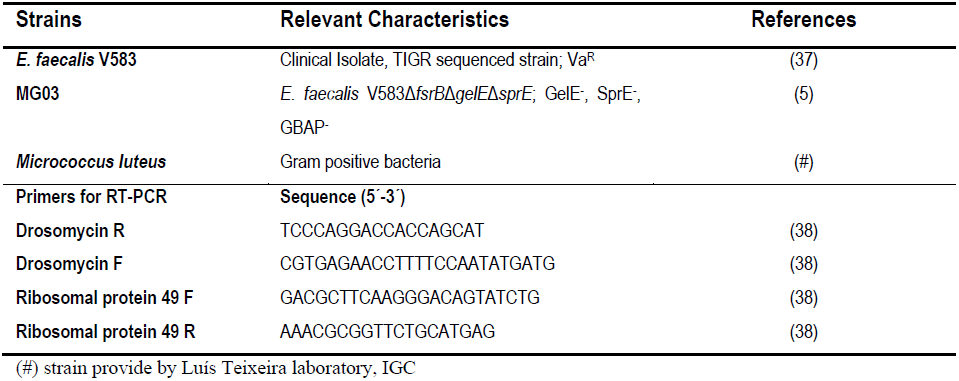
Bacterial strains and primers used in this study.

### RNA extraction and Real-Time PCR for AMP expression

*E. faecalis* and *M. luteus* strains were grown in BHI and LB, respectively, at 37°C, until OD (600nm) 0, 02. The bacterial strains were injected into *W*^1118^ flies. At 6h and 24h after infection 10 flies were collected and homogenized to proceed to RNA extraction. Total RNA extraction was prepared using a TRIzol (Life Technologies) extraction protocol and purified with RNA Clean-up & Concentration from Zymo Research Company. SYBR Green quantitative real-time PCR analysis was performed using 1st Strand cDNA Synthesis kit RT-PCR (AMV) and LightCycler^®^ 96 System from Roche Company. The primers used are listed in Table 1. The amount of mRNA detected was normalized to control rp49 mRNA values. Normalized data were used to quantify the relative levels of a given mRNA according to cycling threshold analysis (ΔCt). Relative ΔCt gene/ΔCt rp49 ratios of unchallenged wild-type controls were anchored in 1 to indicate fold induction. Graphs represent the mean and SD of relative ratios detected in three independent biological repetitions.

### Drosophila

*W*^*1118*^ *Drosophila* male flies (Table 2) were injected with 50 nl of bacteria at OD (600 nm) 0.02 from one of the strains: V583, V583*ΔfsrBΔgelEΔsprE* and *M. luteus*. As control, flies were injected with the same volume of BHI medium. Male flies were anesthetized with CO_2_ and injections were carried out with a pulled glass capillary needle using a nano-injector (Nanoliter 2000, World Precision Instruments). Reproducibility was measured by determining the number of bacteria injected at time zero. Injected flies were placed at 29°C, 65% humidity. Seventy-five flies were assayed for each survival curve, and they were placed in three vials of 25 flies each. Each experiment was repeated three times, making a total of 75 flies tested per strain in each set of three replicates, to ensure high confidence results. Death was recorded at 0, 4, 6, 8, 10, 12, and 24 h hours post-injection. All experiments were performed at least three times. Following challenge with bacteria, six individual flies were collected (at 0 h, 4 h, 8 h, 12 h and 24 h), homogenized, diluted serially, and plated onto Enterococcel agar (Quilaban). *E. faecalis* CFUs (colony forming units) were determined by testing three groups of six flies for each time point.

**Table 2.** Flies used in this study.

| Flies | Relevant Characteristics |
| --- | --- |
| *W*<sup>1118</sup> | Wild type fly |
| *W*<sup>1118</sup>*PPO1*<sup>Δ</sup>*PPO2*<sup>Δ</sup> | Flies without PPO1 and PPO2 (17) |
| *W*<sup>1118</sup>*HmlΔ*>*GFP/UAS* | *W*<sup>1118</sup> flies with hemocytes labeled with GFP |
| *W*<sup>1118</sup>*HmlΔ*>*GFP/UAS-Bax* | *W*<sup>1118</sup> flies without hemocytes |

### *Drosophila* melanization

*Drosophila W^1118^*, in pre-pupa stage, was injected with 50 nl of bacteria at OD (600 nm) 0.02 from one of the strains: V583, V583*ΔfsrBΔgelEΔsprE* and *M. luteus*. As control, flies were injected with the same volume of BHI medium. Injections were carried out with a pulled glass capillary needle using a nano-injector (Nanoliter 2000, World Precision Instruments). The melanization process was recorded at 0, 6, 24 and 48 h hour’s post-injection using the stereoscope Lumar V12 (Zeiss company). All experiments were performed at least three times.

### Statistical analysis

Statistical analysis of *Drosophila* survival was performed using GraphPad Prism software version 5.03. Survival curves were compared using Log-rank and Gehan-Breslow-Wilcoxon tests. Statistical analysis of *Drosophila* survival was performed using Student’s *t-test*.

## RESULTS

In order to understand the mechanisms by which the Fsr-GelE-SprE factors in *E. faecalis* induce fast death of *Drosophila* upon infection, we tested whether known innate immune system pathways are differentially regulated in two *E. faecalis* strains, V583 (wild type) and its isogenic mutant devoid of *fsr, gelE* and *sprE* genes.

### Drosomycin expression is similar during *Drosophila* infection with either V583 or V583*ΔfsrBΔgelEΔsprE* strains

It is known that Gram positive bacteria activate the Toll pathway and that Drosomycin is one of the AMPs produced to kill this group of bacteria (6). One way bacteria use to hamper the immune system of the host is by inhibiting these peptides. Indeed, Park *et al* demonstrated that gelatinase from *E. faecalis* is able to degrade Gm cecropin, an inducible AMP in the insect *Galleria mellonela* (11). We were therefore interested to know whether the presence of Fsr-GelE-SprE influenced the expression levels of AMPs. For that we measured the expression of Drosomycin by qRT-PCR at 6h and 24h post-infection in both V583 and V583Δ*fsrB*Δ*gelE*Δ*sprE* strains and in the control strain *M. luteus*. Interestingly, we found that all strains induced drosomycin expression to similar levels in both time points analyzed, over the period of 24h (Figure 1). These results suggest that the *E. faecalis* virulence factors tested do not regulate AMP production in *Drosophila*.

**Figure 1.**
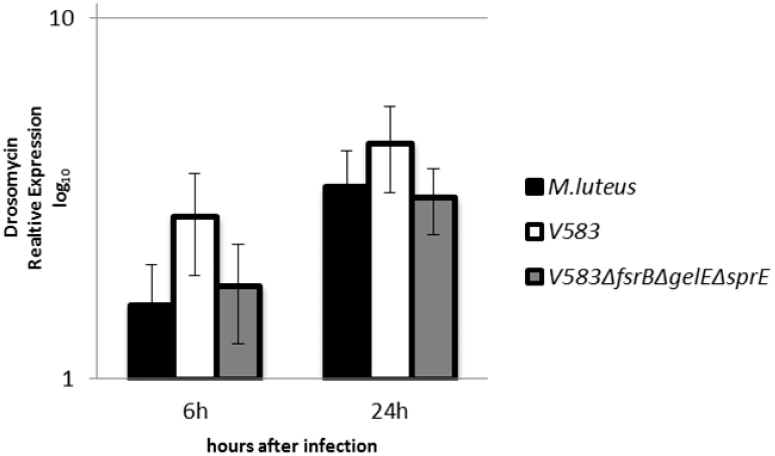
Drosomycin relative expression (scale log10) measured by qRT-PCR. *W*^1118^ flies were challenged by septic injury with Gram positive bacteria: *M. luteus* (black), *E. faecalis* V583 (white) and *E. faecalis* V583Δ*fsrB*Δ*gelE*Δ*sprE* (grey). Total RNAs were extracted at 6h and 24h post-infection. Results were normalized to *rpo49* expression levels. *M. luteus* was used as a positive control of Drosomycin expression and of the Toll pathway activation. Normalized data were used to quantify the relative levels of a drosomycin according to cycling threshold analysis (ΔCt).

### *E. faecalis* Fsr, GelE and SprE do not interfere with *Drosophila* phagocytosis

Phagocytosis is an important defense mechanism that has been conserved during evolution. In *Drosophila* the circulating phagocytic cells are the plasmocytes, which are part of the innate immune system. This complex cellular process is initiated by the recognition of the particles or pathogens to be ingested, followed by cytoskeletal remodeling and signaling events leading to their engulfment and destruction (12). It is known that *E. faecalis* can survive for a prolonged period in mouse peritoneal, human and zebrafish macrophages after being phagocyted (13-15). To investigate whether *E. faecalis* Fsr-GelE-SprE perturb phagocytosis in the fruit fly, we used a *Drosophila* line genetically modified to lack all hemocytes (*W^1118^ HmlΔ>GFP/UAS-Bax*). We found that flies without hemocytes (HmlΔ>GFP/UAS-Bax) show only slightly increased survival rates upon infection with V583 when compared with control flies (W^1118^ HmlΔ>GFP/UAS) (Figure 2A). The same was observed when the two *Drosophila* lines were infected with the *E. faecalis* mutant strain V583*ΔfsrBΔgelEΔsprE* (Figure 2B). The flies died at the same rate with or without hemocytes and regardless of the presence of the *E. faecalis* virulence factors studied. These data suggest that the role of the *E. faecalis* virulence factors tested in host death does not seem to occur through changes in phagocytosis by the hemocytes.

**Figure 2.**
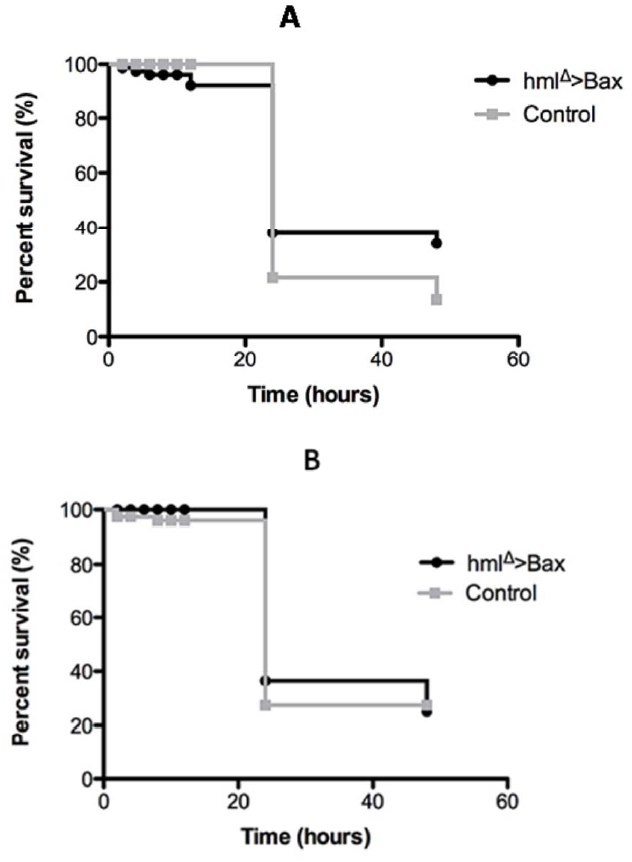
Survival curves of *Drosophila*, with and without phagocytes, infected with *E. faecalis* V583 and *V583*Δ*fsrB*Δ*gelE*Δ*sprE.* (A)*W^1118^ HmlΔ>GFP/UAS-Bax* survival to septic injury with V583wt (B) *W^1118^HmlΔ>GFP/UAS-Bax* survival to septic injury with *V583*Δ*fsrB*Δ*gelE*Δ*sprE*. As a control *Drosophila W^1118^HmlΔ>GFP/UAS* flies were used. For each survival curve, 75 male adult flies, rose at 25°C, where divided in tubes 25 flies each, and infected, by septic injury onto the thorax with thin needle. Data is representative of three independent experiments. Statistical analysis of *Drosophila* survival was performed using GraphPad Prism software version 5.03. Survival curves were compared using Log-rank and Gehan-Breslow-Wilcoxon tests and they were not statistically different.

### Fsr-GelE-SprE leads to increased melanization in pre-pupae

One of the key immune reactions in *Drosophila* is the activation of tyrosinase-type phenoloxidases (POs), which catalyze several reactions leading to the crosslinking of proteins, the production of reactive intermediates with potential cytotoxic activity and ultimately to the production of melanin (16). Melanization is the earliest reaction against the evasion of pathogen and it is visible by the blackening of wound site. To determine if melanization is affected by the presence of Fsr-GelE-SprE in infecting *E. faecalis*, we injected wild type pre-pupae, a stage that allows the easy detection of melanized dark spots, with *E. faecalis* V583 and *E. faecalis* V583Δ*fsrB*Δ*gelE*Δ*sprE* strains. At 6h post-infection, melanized spots are only around the site of injection in all strains analyzed (Figure 3).

**Figure 3.**
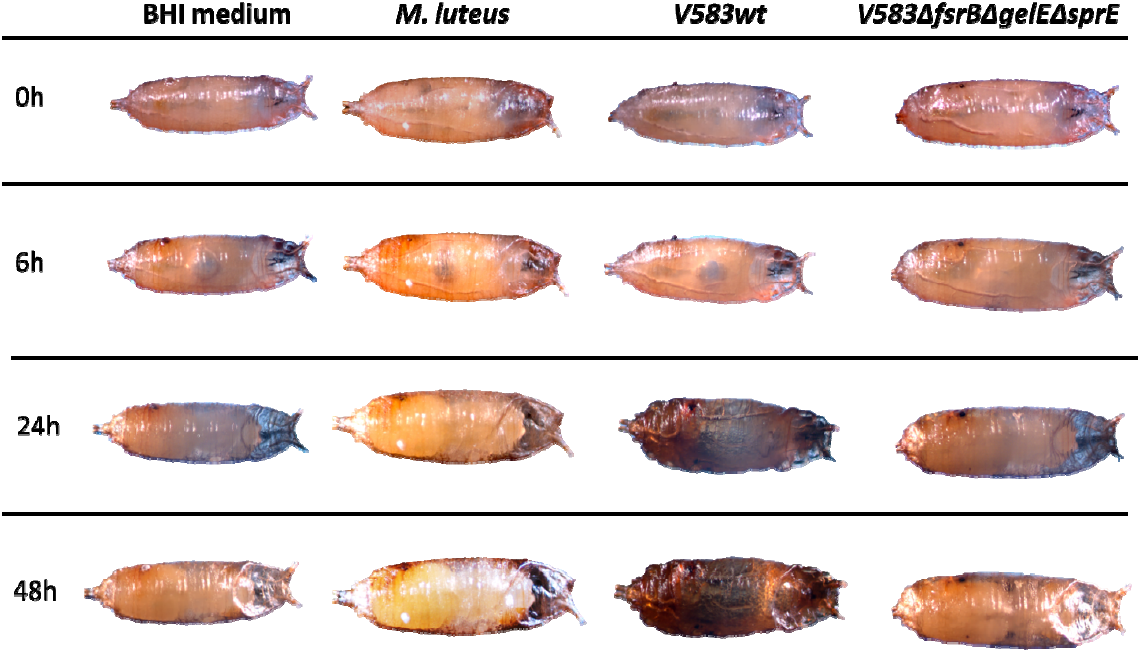
Melanization in wild type fly pre-pupae after infection. *W*^1118^ pre-pupae were infected by septic injury with 50nl of *M. luteus* at 0, 02 OD; V583wt and V583Δ*fsrB*Δ*gelE*Δ*sprE* at 0, 02 OD, and placed at 29°C. Injection with BHI medium and *M. luteus* are controls of this experiment. Pictures were taken with stereoscope Zeiss Lumar V12 after 0h, 6h, 24h and 48h post-infection. This procedure was made at least in 10 pre-pupae and the results were always the same. After 24h hours an exacerbated melanization in the pre-pupae infected with V583wt was observed. All the other pre-pupae showed only the normal black dots around the injection site.

After 24h it is clear that the pre-pupae infected with V583 strain have an exacerbated melanization, which is observed all over the body. In pre-pupae infected with *E. faecalis* triple mutant, however, melanization remains restricted to the wound site, similar to pre-pupae infected with the *M. luteus* control strain. Moreover, pre-pupae infected with wild type bacteria were dead after 24h whereas those infected with the mutant bacteria were still alive after 48h. These results indicate that the presence of the Fsr-GelE-SprE *E. faecalis* virulence factors interferes with the melanization process during infection through which it contributes to host death.

### *E. faecalis* virulence factors modulate melanization effect on *Drosophila* survival

We thus asked if the excessive melanization was responsible for the fast and massive death of the infected hosts. To answer this, we infected flies mutated in two prophenoloxidases (PPO1 and PPO2), which makes them unable to produce melanin (9). Figure 4 shows the survival rates of *W^1118^PPO1^Δ^PPO2^Δ^* mutant and wild type flies infected with V583 and mutant strains. When we compare PPO mutant and control flies infected with the same wild type bacteria, survival rates are similar. However, they have different shapes. During the first 12h of infection *PPO1^Δ^, PPO2^Δ^* flies were affected in their capacity to survive infection by V583 strain while the bacterial counts were slightly higher during this same period, compared to wild type *Drosophila*. Similar survival behavior was observed in *PPO1^Δ^, PPO2^Δ^* flies infected with OG1-RF *E. faecalis* strain (17), which carries the same Fsr-GelE-SprE virulence factors (18). This decreased survival was not observed when *PPO1^Δ^ PPO2^Δ^* flies were infected with the bacterial mutant strain (Figure 4B). Taken together, these results (Figures 4 and 5) suggest that melanization may be involved both in resistance and tolerance to *E. faecalis*, and that its role depends on the presence of the Fsr-GelE-SprE virulence factors. The two bacterial proteases present in V583, GelE and SprE, are able to degrade host structural proteins, thus causing tissue damage, which must be healed in order for the fly to maintain its healthy status. In the absence of melanization, which contributes to tissue healing, it is possible that the flies tolerate less the presence of V583 carrying Fsr-GelE-SprE factors.

**Figure 4.**
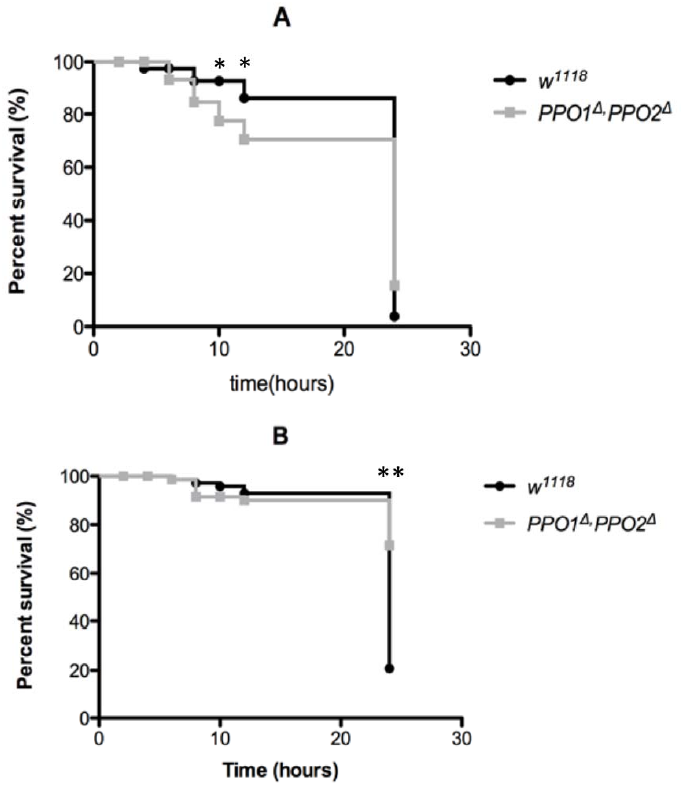
Survival curves of *Drosophila* with and without melanization. *Drosophila* W^1118^ and *PPO1*^Δ^*PPO2^Δ^* survival to septic injury, with V583wt (A); and V583Δ*fsrB*Δ*gelE*Δ*sprE* (B) strains. For each survival curve, 75 male adult flies, rose at 25°C, where divided in tubes 25 flies each, and infected, by septic injury onto the thorax with a thin needle. Data is representative of three independent experiments. Statistical analysis of *Drosophila* survival was performed using GraphPad Prism software version 5.03. Survival curves were compared using Log-rank and Gehan-Breslow-Wilcoxon tests and they were not statistically different. Survival rates at time point 12h and 24h are marked with (*) to represent statistically different results (calculated using the Student’s *t-test*) from the respective wild-type (*p < 0.05; **p < 0.005).

**Figure 5.**
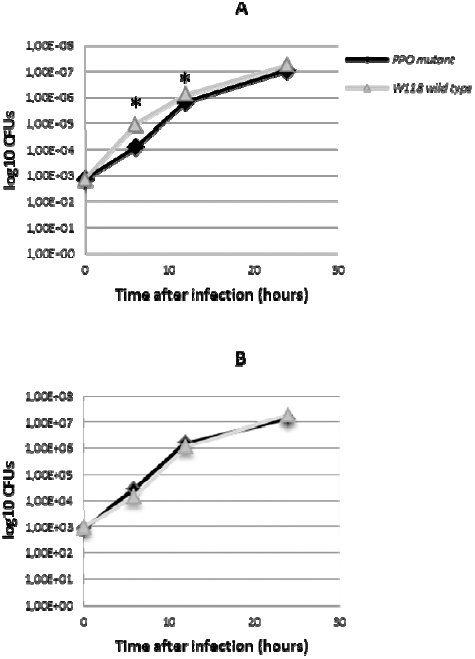
*E. faecalis* growth curves in injected flies. (A) *E. faecalis* V583 growth rates in injected fly W^1118^ and *W^1118^PPO1^Δ^PPO2^Δ^*. (B) *E. faecalis* V583*ΔfsrBΔgelEΔsprE* growth rates in injected fly W^1118^ and *W^1118^PPO1^Δ^PPO2^Δ^* Male adult flies (5- to 7-day-old), raised at 25°C, were divided in tubes of 25 flies each, and infected, by septic injury onto the thorax with a thin needle, with V583 strains. Flies were collected at 0, 6, 12, and 24 h. Three groups of six flies for each time point were homogenized and plated in Enterococcel agar and *E. faecalis* CFUs were determined. Student’s t-test was used for statistical analysis. Asterisks (*) indicate the statistical significance (*p < 0.05; **p < 0.005).

When we compare the survival rates of PPO mutant and control flies infected with the mutant bacteria major differences were observed. 24 hours after infection, 80% of the *PPO1^Δ^PPO2^Δ^* mutant flies infected with V583Δ*fsrB*Δ*gelE*Δ*sprE* triple mutant were still alive (Figure 4B). In contrast, all wild type flies were dead when infected with the wild type bacteria (Figure 4A). These results clearly indicate that when both melanization and Fsr-GelE-SprE are absent almost all flies survive infection.

## DISCUSSION

The fly mechanisms responsible for protection against bacterial infections are not clearly understood yet. *Drosophila* has four distinct pathways implicated in regulation of genes induced upon septic injury, namely Toll, Imd, JNK and JAK-STAT (19). Previous studies have shown that *E. faecalis* induces both cellular and humoral immune response mechanisms in *Drosophila*. Toll seems to be the crucial pathway in the defense against *E. faecalis* (20): whereas Toll pathway mutants are susceptible to *E. faecalis*, Imd mutants are not (19). The Toll pathway is responsible for production of several AMPs: diptericin, cecropin, drosocin and attacin are active against Gram-negative bacteria and drosomycin, metchnikowin and defensin to fungi and Gram-positive bacteria. Except for defensin, *E. faecalis* is resistant to the bactericidal activity of all AMPs produced by *Drosophila*, and even from *G. mellonella* (21). It is thus not clear how the Toll pathway confers protection against Gram-positive bacteria, as it is known that defensin is not necessary to mediate protection (20). Previous *in vitro* studies showing that the proteases GelE and/or SprE may degrade insect AMPs, have led researchers suggest that *E. faecalis* success in insect species could be attributed to the degradation of the host innate immune AMPs by the proteases. However, in a previous study, our findings suggest otherwise (5). In fact, as we observed no difference in growth inside the host between any of the mutants and wild type V583, we conclude that neither the Fsr system nor the proteases it controls affect bactericidal action by the fly. This implies that none of the proteases provides self-protection against any AMP in the fly immune system (5). In the present study we showed that the presence of Fsr-GelE-SprE does not affect the levels of drosomycin expression, further supporting the likely irrelevant role of *Drosophila* AMPs on *E. faecalis* infection progression. However, this does not exclude the possibility that the presence of these proteases in high amounts may turn the host more fragile to other bacteria due to AMPs degradation. In fact, previous work has shown that GelE is able to degrade host AMP´s and that this is responsible for insects getting less able to deal with *Escherichia coli* strains (11).

The way the fly is able to fight invading microorganisms also includes a cellular immune response that can result in the phagocytosis of relatively small organisms like bacteria or the encapsulation of larger parasites (22). However, little is known about how *Drosophila* phagocytes affect the course of infections (23). On the other hand, bacteria that are specialized in growing inside phagocytes have developed ways to fight these cells from within. Moreover, previous studies have demonstrated that the pathogenesis mechanisms developed by *Mycobacterium marinum* and *Listeria monocytogenes* to fight vertebrate phagocytes also function in the fly (24, 25). In the case of extracellular pathogens, such as *E. faecalis*, it is known that these bacteria are able to stand macrophages defense mechanisms for hours and days (26). Although some *E. faecalis* defense mechanisms have been implicated in its prolonged life inside macrophages (27-29), neither Fsr nor the two proteases it regulates seem to play a role in bacterial survival inside these defensive cells. Recently, macrophages in zebrafish were shown to phagocytize bacteria in blood circulation being only able to engulf surface-associated microbes (15). It is also known that homolog of tumor necrosis factor (TNF) encoded by *eiger* is required for innate immune responses that are effective at fighting extracellular pathogens but are wasteful or simply ineffective when fighting intracellular pathogens (30). In our model, despite being phagocytized by *Drosophila* hemocytes (results not shown), neither Fsr nor GelE or SprE were found to affect the cellular immune response of *Drosophila*.

Melanization is another *Drosophila* immune response. It is visible by the blackening of a wound site or the surface of pathogens, which results from the synthesis and deposition of melanin. In addition to being important for wound healing, melanin can encapsulate and sequester pathogens, and the reaction intermediates appear to be directly toxic to microbes as well (31). In our study, the effect of the presence of *E. faecalis* virulence factors Fsr-GelE-SprE in melanization was evaluated. Pre-pupa infected with *E. faecalis* V583 strain showed a different melanization pattern from that shown by pre-pupa infected with the triple mutant strain. Consistent with the excessive and all over body melanization observed with V583 infection, pre-pupa died earlier with this virulent strain. The phenotype of these pre-pupa at 24h post-infection resembles that of pupa devoid of serpin27 and PPO2 proteins (17), which present high levels of constitutive PO activity. Activation of melanization is strictly regulated. Uncontrolled melanization generates excessive toxic intermediates that can kill the host (9). Recognition of pathogens and injury leads to the activation of a serine protease cascade that culminates in proteolytic cleavage of inactive PPO to active PO. Serine protease inhibitors, called serpins, are responsible for keeping the melanization strictly localized at the site of injury or infection (9). Manipulation of the PO activity, through interfering with the proteolytic activation of the melanization cascade, is a strategy developed by some pathogens (32). The bacterial proteases studied GelE and SprE, are known to degrade host proteins and cause tissue injury, and although their ability to interfere with the PO activation was never evaluated. Our pre-pupa results suggest one of two hypotheses: that the *E. faecalis* virulence factors are involved in melanization de-repression and massive release of PO activity from crystal cells in the hemolymph; or that the proteases induce such a high degree of tissue injury that the massive activation of the melanization for tissue healing becomes overwhelming and deleterious to the host itself.

If the first hypothesis was to be true, infection of *Drosophila PPO1^Δ^, PPO2^Δ^* mutant with V583, which carries the proteases, would result in reduced host death. However, the proteases, and the Fsr system that regulates their expression, were found not to play a part in death by melanization in adult flies. A possible explanation for these results could be related with a different role and impact of crystal cells in pre-pupae and adults. As reported by Binggeli *et al* (2014), crystal cells could have evolved as an adaptation to release a large quantity of PO activity in the hemolymph of pupa(17). Therefore, a role for the bacterial proteases in induced PO activity, which would cause host death, cannot be ruled out in the pre-pupa developmental stage, and is likely through activation of melanization through massive tissue injury.

In adult flies, infection with the V583 mutant did not lead to increased survival. This was unexpected as we previously reported that Oregon flies showed increased tolerance to the same V583 mutant (5). This result shows that, within the same host species, the host genetic background plays an important role in the outcome of an infection by an opportunistic pathogen, such as *E. faecalis*. The difficulty in studying *E. faecalis* virulence is, once again, evidenced with this result. Interestingly, the host genetic background is not the only caveat of *E. faecalis* virulence studies. The bacterial genomic content is also important. In fact, the *PPO1^Δ^, PPO2^Δ^* mutation had a different impact in survival of the flies when infected with the two *E. faecalis* bacterial strains. *PPO1^Δ^, PPO2^Δ^* mutant flies were more tolerant to *E. faecalis* deprived of its major virulence factors than the wild type *Drosophila*. The presence of the proteases completely altered the PO activation impact on *Drosophila* tolerance to *E. faecalis*. The current knowledge on *Drosophila* immunity and on *E. faecalis* virulence, and on their interaction as host and pathogen, is not enough to provide solid explanation for this finding. Although knowledge of the effects of having no PO activity is scarce, some reports have highlighted the facts that affecting one particular immune pathway, leads to changes in other *Drosophila* functions. The *Drosophila* mutant strain used in this study has been partially, and recently, characterized and found to show increased Toll activation by Gram-positive bacteria, when compared to its wild type (17). Activation of Toll leads to induction of immunity, but also to reallocation of host resources, by suppressing insulin signaling throughout the organism, leading to a decrease in both nutrient stores and host growth (33). Immune and metabolic rearrangements in the *Drosophila* PPO mutant flies may be the cause of the observed increased tolerance to infection by the *E. faecalis* mutant strain. Future studies should clarify this.

In humans melanization does not occur, however PPO activation mediated by a serine protease cascade is somewhat analogous to the coagulation pathway and complement system (CS) in human plasma (34). Like the melanization, the fast activation of the complement system after a microorganism infection of a potential host is a crucial step in clearance of many pathogens. For example, anaphylatoxins like C3a and C5a, products of the CS cascade, are commonly involved in exacerbated inflammatory reactions that can cause direct harm to the host following infections (35). We know that GelE destroys the C3a complement of human cells and AMPs of *G. mellonella* (11, 36). Taking into account that the serine protease cascade during melanization is analogous to the complement system, we hypothesize that in humans, Fsr regulated components interfere with the complement response. Future studies should investigate this hypothesis.

Our study shows that the outcome of *Drosophila* infection with *E. faecalis* depends both on the bacteria gene content (in particular, the presence of Fsr and the proteases it regulates) and on the host immunity status (in particular, on the PO activity). This stresses the need to know more about how the host reaction is altered when the pathogen changes. As pointed out by others (10), a systematic genome-wide exploration of pathogen mutants and their interaction with fruit fly immunity is important. The present study points out some interesting facts that should orient future studies aiming at finding new ways to diminish the mortality associated with *E. faecalis* infections. It may not be sufficient to shut off the bacterial virulence, namely through interference with the quorum-communication. An efficient control of *E. faecalis* infection outcome should also include the host immune manipulation. The conservation between the innate immune system of humans and *Drosophila* will allow future studies to develop new targets to control *E. faecalis* infections in humans.

## ACKNOWLEDGEMENTS

The work was supported by FCT through grant #Pest-OE/EQB/LAO004/2011. Neuza Teixeira was supported by FCT fellowship SFRH/BD/65750/2009. The authors are grateful to Bruno Lemaitre from Global Health Institute, School of Life Sciences, École Polytechnique Fédérale Lausanne (EPFL)- Switzerland for the *W^1118^ PPO1^Δ^PPO2^Δ^ Drosophila* lines; and to Luis Teixeira from Instituto Gulbenkian de Ciência, Oeiras – Portugal for supplying the *M. luteus* strain. We are also grateful to Anabela Bensimon-Brito for the technical support in qRT-PCR experiment, to Carolina Moreira and Ana Sofia Brandão for the technical support in *Drosophila* phagocytosis experiments, and to Lara Carvalho for comments and revision of the manuscript.

